# Phylogenetic analysis by SNP and development of SSR marker in *Passiflora*

**DOI:** 10.1101/2020.07.15.203513

**Authors:** Yanyan Wu, Qinglan Tian, Weihua Huang, Jieyun Liu, Xiuzhong Xia, Xinghai Yang, Haifei Mou

## Abstract

Information of the *Passiflora* genome is still very limited. Understand the evolutionary relationship between different species of *Passiflora*, and develop a large number of SSR markers to provide a basis for the genetic improvement of *Passiflora*. Applying restriction site associated DNA sequencing (RAD-Seq) technology, we studied the phylogeny, simple sequence repeat (SSR) and marker transferability of 10 accessions of 6 species of *Passiflora*. Taking the partial assembly sequence of accessions P4 as the reference genome, we constructed the phylogenetic tree using the detected 46,451 high-quality single nucleotide polymorphisms (SNPs), showing that P6, P7, P8 and P9 were a single one while P5 and P10 were clustered together, and P1, P2, P3 and P4 were closer in genetic relationship. Using P8 as the reference genome, a total of 12,452 high-quality SNPs were used to construct phylogenetic tree. P3, P4, P7, P8, P9 and P10 were all single branch while P1 and P2 were clustered together, and P5 and P6 were clustered into one branch. A principal component analysis (PCA) revealed a similar population structure, which four cultivated passion fruits forming a tight cluster. A total of 2,614 SSRs were identified in the genome of 10 *Passiflora* accessions. The core motifs were AT, GA, AAG etc., 2-6 bases, 4-16 repeats, and 2,515 pairs of SSR primer were successfully developed. **T**the SSR transferability in cultivated passion fruits is the best. These results will contribute to the study of genomics and molecular genetics in passion fruit.

## 1. Introduction

*Passiflora* is an herbaceous plant or vine, consisting of 16 genera and 530 species[1]. At the present, the cultivated passion fruits are mainly purple passion fruit (*Passiflora edulis* Sims) and yellow passion fruit (*Passiflora edulis* var. *flavicarpa* Degener). Passion fruits are popular for consumers because of their aromatic aroma and richness in sugar, vitamins and mineral elements[2,3].

While Approximately 360 plants have been sequenced (http://www.plabipd.de/index.ep), there is still no reference genome in passion fruit. Our cognition of the *Passiflora* genome is still very limited. Santos *et al*. used the bacterial artificial chromosome (BAC)-end sequencing method to study the *Passiflora* genome and found that it was rich in repetitive elements, among which the number of reverse transposons (Class I) was very abundant[4]. Araya *et al*. sequenced the cultivated passion fruit CGPA1 with Illumina GAII sequencing and analyzed 28 *Passiflora* accessions using SSR markers. They found that the genetic clustering of *Passiflora* was not related to geographical origin[5]. Munhoz *et al*. identified and annotated the repetitive sequences in the BAC-end library and assembly partial sequence of *Passiflora*. Subsequently, they conducted microsyntenic region analysis with *Populus trichocarpa* and *Manihot esculenta* of Malpighiales species, and found that *Passiflora edulis* gene-rich segments were more than those the other two. The fragment of *Passiflora edulis* is more compact, even though its genome is larger [1].

SSR markers are characterized by high polymorphism, good reproducibility, co-dominance and low amount of DNA required for detection. Especially, they can be used in discrimination for genetic differences in accessions with closer genetic relatives. Oliveir *et al*. [6] identified the first microsatellite loci from the *Passiflora* genome microsatellite enrichment library, and developed 10 pairs of SSR primers to verify them with 12 yellow passion fruits, among which, 7 pairs of SSR primers were polymorphic. Padua *et al*. [7] screened out 53 positive clones, identified 8 SSRs, and designed 7 pairs of primers to genotype 12 population groups. They found that these SSR markers had very high frequency of polymorphism. Subsequently, Cerqueira-silva *et al*. [8]also developed 25 pairs of SSR primers using this method, of which, 7 pairs were polymorphic. Cazé *et al*. [9] identified SSR using the *Passiflora* genome microsatellite enrichment library, designed 12 pairs of SSR primers, and verified the 7 pairs with polymorphism using two wild species of *Passiflora contracta*. In 2014, Cerqueira-silva *et al*. [10]developed 69 pairs of SSR primers using two *Passiflora* genome microsatellite-enriched libraries and found that 43 pairs of primers were polymorphic. Silva *et al*. [11] used 31 pairs of SSR primers to study the transferability of SSR markers between wild and cultivated passion fruits, and found that a part of these SSR markers exhibited very high transferability (71.4%), indicating that it is feasible to establish the SSR transferability between the wild and cultivated passion fruits. In 2017, Araya *et al*. [5]developed 816 pairs of SSR primers using partial sequences of the cultivated passion fruits genome sequencing, validated them in 79 wild and cultivated passion fruits and found that 42 out of 57 pairs of SSR primers were polymorphic. da Costa *et al*. developed 42 pairs of SSR primers using cDNA sequences in a suppression subtractive hybridization library and found that 34 pairs of primers between *Passiflora edulis* and *Passiflora alata* have high transferability rates [12]. However, to date, the number of effective SSR primers developed and validated by researchers is still limited, and it is still unable to meet the need for research on the molecular genetic map, gene mapping and marker-assisted selection for*Passiflora*.

In this study, we sequenced 10 *Passiflora* accessions using RAD-Seq. Our study aimd (1) to evaluate the enzymatic cleavage effect of restriction endonucleases EcoRI in different *Passiflora* accessions, (2) to construct the phylogenetic relationship between the cultivated passion fruitsand its related species using SNPs, and (3) to develop a large number of SSR markers derived from different *Passiflora* species.

## 2. Materials and Methods

### 2.1 Materials and Planting

Ten *Passiflora* accessions were as follows:Jinlingxinxinguo (*Passiflora edulis* Sims), Baleweihuangjinguo (*Passiflora edulis* var. *flavicarpa* Degener), Yunanhuangguoyuanshengzhong (*Passiflora edulis* var. *flavicarpa* Degener), Ziguoqihao (*Passiflora edulis* Sims), Lvpibaixiangguo (*Passiflora edulis* Sims), Languan passion fruit (*Passiflora caerulea* Lim), Honghua passion fruit (*Passiflora coccinea*), Banna passion fruit (*Passiflora xishuangbannaensis*), Daguo passion fruit (*Passiflora quadragularis* Linn), and Sweet granadilla (*Passiflora ligularis*),they were numbered P1-P10 in turn. *Passiflora xishuangbannaensis* is a new species discovered in 2005 and is currently found only in the wilderness area of Yunnan Province, China [13]. These ten *Passiflora* accessions were planted in the germplasm resource nursery (22.85° N, 108.26° E) of the Biotechnology Research Institute, Guangxi Academy of Agricultural Sciences in 2017.

### 2.2 DNA Extraction and Illumina Sequencing

The young leaves of 10 *Passiflora* accessions in the 6-8 leaf stage were collected and store in a -20 °C refrigerator. Genomic DNA was extracted by cetyl trimethylammonium bromide (CTAB) method [14]. The purity and integrity of each sample DNA were examined by agarose gel electrophoresis. Nanodrop (Thermo Fisher Scientific, Waltham, MA, USA) was used to detect the purity of DNA with OD260/280 being equal or higher than 1.8. Qubit (Thermo Fisher Scientific) was used to accurately quantify the DNA concentration. The lowest concentration was 50 ng/μl.

The RAD library was constructed by reference to the method described by Baird *et al*. [15]. The qualified DNA samples were digested with EcoRI, randomly fragmented, and the entire library was prepared by terminal repair, addition of a tail, addition of sequencing linker, purification, and PCR amplification. The constructed library was sequenced by Illumina NovaSeq 6000.

### 2.3 Assembly of Reference Genome

Until now, there has been no *Passiflora* reference genome and a partial genome needs to be assembled for SNP analysis. To evaluate the quality of the sequencing data, CD-HIT-EST[16]was used to cluster the high-quality reads, and then the Velvet optimization program Velvetopt[17]was used to locally assemble each of the selected samples to obtain the final assembly sequences.

### 2.4 SNP Detection

SNP detection: (I) Using the BWA alignment software, the PE reads of 10 clean data sets of *Passiflora* sets were compared with the reference genome; (II) the alignment results were converted into SAM/BAM files using SAMtools; (III) the alignment ratio and coverage were compared with Perl scripts; and (IV) the results were compared for SNP detection using SAMtools.

### 2.5 Phylogenetic Tree, Population Sturcture and Principal Component Analysis

The neighbor joining (NJ) phylogenetic tree was built using MEGA7 software. The number of Bootstrap repetitions was 1000, and other parameters were default. The population structure was analysed by ADMIXTURE software[18], which the subpanel number was predicted from 1 to 8. Principal component analysis (PCA) was performed using GAPIT[19].

### 2.6 SSR Primer Design and PCR Analysis

SSR Search was used for SSR detection. The specific parameters were set as follows: the minimum length of the SSR repeating unit was 2 and the maximum length of the repeating unit was 6, the minimum length was 12, the lengths of the upstream and downstream sequences were 100 bp, and the minimum SSR distance was 12 bp, and then the SSR search of the cultivated passion fruit genome sequencing was carried out to obtain the genome sequence containing SSR.

According to the SSR flanking sequence, the Primer3 was used to design primers with the optimal length of 24 bp, the shortest length of 20 bp, and the longest length of 28 bp. The optimum annealing temperature was 55 °C, the lowest annealing temperature was 53 °C, and the highest annealing temperature was 58 °C. The maximum value of the annealing temperature difference was 1 °C.

PCR amplification was conducted using a 15 μl reaction system consisting of 1 μl of primer, 2 μl of template DNA, 7 μl of Instant PCR Kit 3.0 PCR Mix (Tiandz, Beijing, China), and 5 μl of double distilled water to final volume. The reaction procedures were set as follows: pre-denaturation at 94 °C for 3 min, 35 cycles of 94 °C for 45 s, 55 °C for 30 s, 72 °C for 30 s, and finally 72 °C extension for 5 min. The annealing temperature and the extension time were adjusted depending on the size of the target product, respectively. The PCR product was tested on a 6% polyacrylamide gel electrophoresis gel.

## 3. Results

### 3.1 Rad-Seq Data Analysis

Ten *Passiflora* accessions were sequenced to gain a total of 14.68 Gb raw data, and the filtered clean data was 14.586 Gb. The raw data of each sample was between 1.15 Gb and 1.83 Gb, and the sequencing quality was quite high (Q30 >= 93.2 %), the GC content was between 39.6% and 41.9%, and the ratio of the number of enzyme-captured reads to the number of reads after de-weighting was between 97.1% and 99.0% (Table 1). Therefore, the data volume of all samples was sufficient, the sequencing quality was qualified, the GC distribution was normal, and the database was successfully sequenced, and thus, subsequent analysis could be performed.

The CD-HIT-EST was used to cluster the samples containing the enzyme recognition sites. The filtering criteria were between 10-400 for each type of reads in the class and the cut tag numbers were between 13,725 and 81,605 for the number of filtered classes. The cut pair reads were 332,415-1,375, and 313, and the cut/pair ratios were in the range of 10.7% - 34.3% (Table 2). We selected the cut/pair two extreme values of P4 and P8 for sequence assembly and performed SNP analysis as a reference genome.

### 3.2 SNP Detection

Burrows-Wheeler Aligner (BWA) was used to compare the clean data of 10 samples with the P4 genome. The comparison results used SAMtools for sort and rmdup, and the BAM files after sorting and de-duplication were counted. The mapped reads were 2118474-6708533, and the mapping_rate was 19.2%-61.5%. Average depth was 96.9-139.5×, Coverage 1X was 40.1%-99.0%, and Coverage 4X was 18.1%-91.9% (Table 3).

We used BWA to compare the clean data of 10 samples with the P8 partial genome. The results were compared using SAMtools for sort and rmdup. The BAM files were sorted and de-duplicated. The mapped reads were 932975-8528271 and the mapping rate was 7.9%-78.0. %, Average depth was 31.6-113.1×, Coverage 1X was 3.9%-99.4%, and Coverage 4X was 1.9%-86.9%% (Table 4).

SNP mainly refers to DNA sequence polymorphism caused by variation of a single nucleotide at the genome level, including single base conversion, transversion and the like. SNP detection was performed on 10 samples using SAMtools. After filtration, high quality SNPs were finally obtained for subsequent analysis. A total of 46,451 and 12,452high-quality SNPs were obtained using P4 and P8 as a reference gene, respectively(Table 5).

### 3.3 Phylogenetic Tree, Population Structure and PCA Nanlysis

Using P4 as the reference genome, the phylogenetic trees of 10 *Passiflora* accessions were constructed using 46,451 high-quality SNPs. The accessions P6, P7, P8 and P9 alone were all one branch while P5 and P10 were clustered together, and P1, P2, P3 and P4 were closer in genetic relationship (Figure 1a).

**Figure 1.**
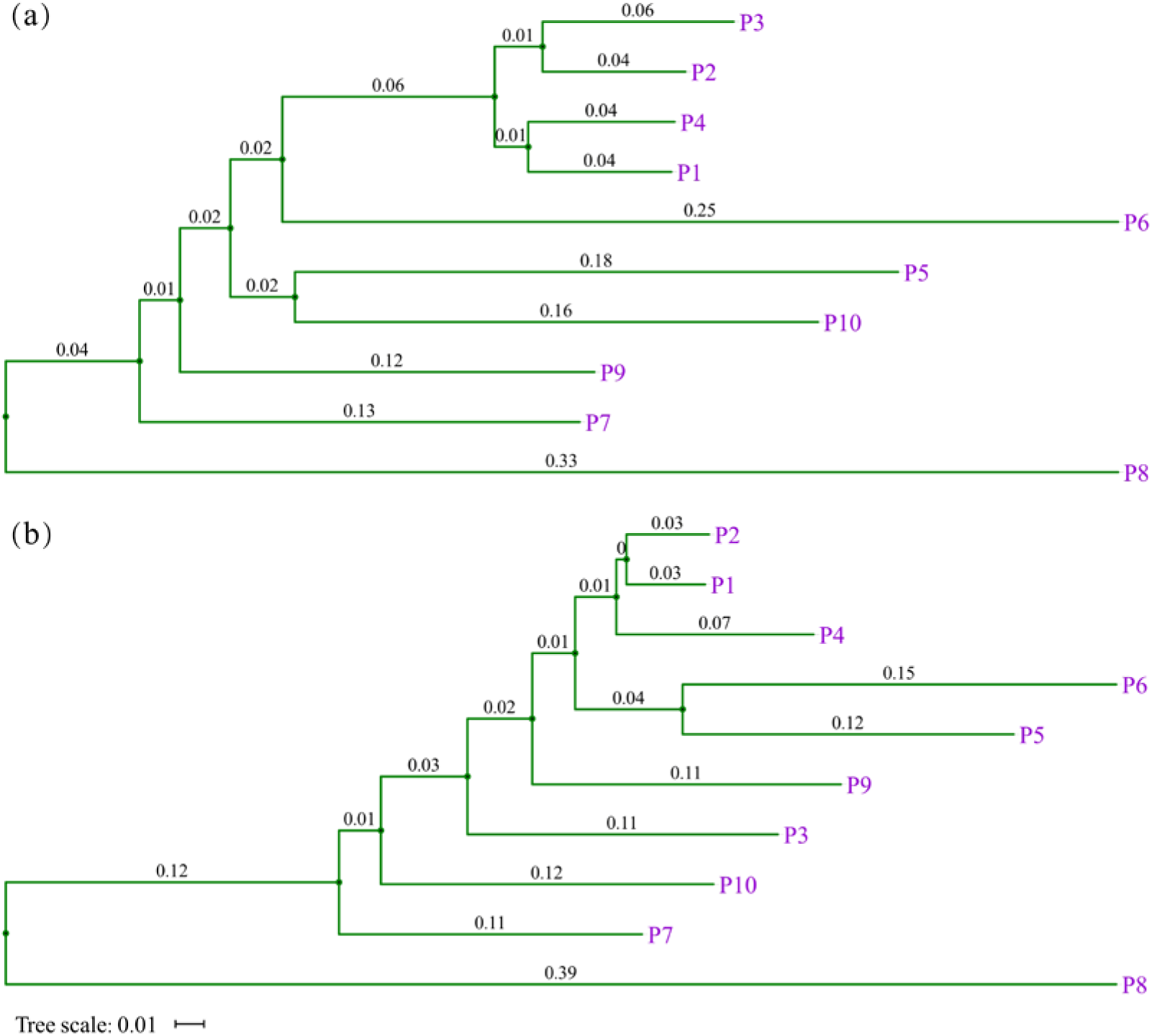
The phylogenetic trees of 10 *Passiflora* accessions. (a) P4 as the reference genome; (b) P8 as the reference genome.

Using P8 as the reference genome, the phylogenetic trees of 10 *Passiflora* accessions were constructed using 12,452 high-quality SNPs. *Passiflora* accessions P3, P4, P7, P8, P9, and P10 individually were all clustered in one branch each while P1 and P2 were clustered together, and P5 and P6 were grouped together (Figure 1b). The phylogenetic trees of 10 *Passiflora* accessions constructed with P4 and P8 as reference genomes were partially different.

Based on 46,451 high-quality SNPs, the population structure analysis indicated that 10 accessions were divided into three classes (Fig 2a,b). The result of PCA analysis were consistent with the population structure analysis. Four cultivated passion fruits were classified into one subpopulation, and P8 was a separate category (Fig 2c).

**Figure 2.**
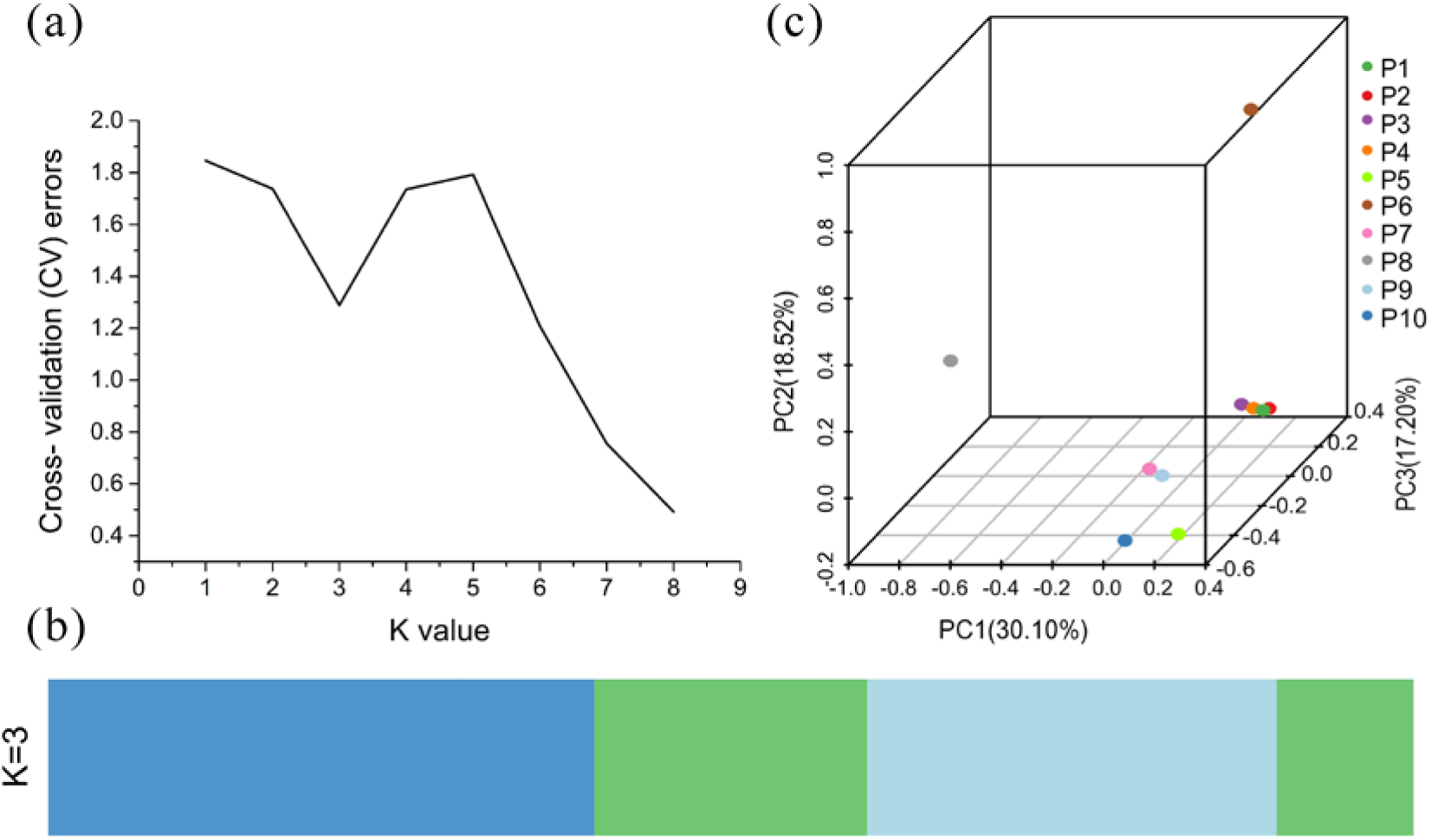
Population structure and PCA analysis of 10 *Passiflora* accessions. (a,b) Phylogenetic relationships based on the population structure, presenting the distribution of K = 3 genetic clusters with the smallest cross-validation error, (c) Principal components for the variation among accessions.

### 3.4 SSR Detection and Primer Development

The other 8 *Passiflora* accessions were sequenced and assembled by the same method to obtain the final assembly sequences, and the contig below 100 bp were filtered through. The results of partial assembly are shown in Supplementary Table 1. Total contig number were in range of 6,964-71,090, average contig length was 282-353 bp, N50 length was 335-403 bp, accounting for 39.43%-42.54%.

Subsequently, SSR search was used to perform SSR detection on the assembled 10 genome sequences of *Passiflora*. The number of SSRs varied greatly among the varieties, and 755 were detected in P8 while only 16 in P9. A total of 2,614 SSRs were identified, the number of bases of SSR core motifs were 2-6, the core motifs of SSR were AT, GA, AAG, etc., and the number of SSR repetitions ranged 4-16.

The identified flanking sequences derived from the SSRs in the cultivar genome were 100 bp each. Next, the primer design software Primer3 was used to design primers for the filtered SSR markers, and a total of 2515 pairs of primers (Supplementary Table 2) were successfully developed.

### 3.5 SSR Marker Validation and Transferability Assessment

In the above-mentioned developed SSR markers, 5 pairs of primers were randomly selected for each accession, and a total of 50 pairs of primers (Table 6) were used to validate the transferability of 10 *Passiflora* accessions. All 50 pairs of primers were able to amplify the target band in 10 accessions. The transferabilities were in the range of 0-100% with the average transferability being 54.22% (Supplementary Figure 1). Pe1001-Pe1005 was derived from the cultivated passion fruit, which can amplify the target band in other species of *Passiflora* as well.

In *Passiflora caerulea*, only the marker Pe10028 was able to amplify the target band in Lvpi passion fruit, the transferability was 2.22%. None of the five SSR markers of *Passiflora xishuangbannaensis* could amplify the target band in the other nine species of *Passiflora*.

## 4. Discussion

Reduced-representation genome sequencing (RRGS) requires restriction enzyme digestion of genomic DNA, an important parameter to measure whether the enzymatic digestion effect is good and whether the library is qualified. In this study, we used restriction endonuclease EcoRI (recognition sequence▾GAATTC) to digest ten *Passiflora* accessions with a cut rate of ≥97.12%. Therefore, EcoRI can be used as a n endonuclease for RRGS of *Passiflora*.

As the third generation of molecular markers, SNP has many characteristics, including uniform distribution, rich polymorphism and high precision[20]. With the improvement of high-throughput sequencing technology, cost reduction and rapid development of bioinformatics, a large number of SNPs have been identified[21]. In the present study, the number of SNPs identified in different species of *Passiflora* as reference genomes was significantly different. The genetic evolution analysis of 10 *Passiflora* accessions using two different numbers of SNP showed the same results. *Passiflora xishuangbannaensis* was distantly related to the other 9 *Passiflora* accessions. Banna passion fruit is a Chinese wild passion fruit[13], which is far from the South American origin of *Passiflora*. These results suggest that there is a high genetic diversity among the geographically different germplasms.

Among the five cultivated passion fruits, either P4 or P8 is reference genome, Jinlingziguo, Baleweihuangjinguo, Yunnanhuangguoyuanshengzhong, Ziguoqihao, the four *Passiflora* accessions are closer in genetic relationship. Lvpi passion fruitmay be a cultivated passion fruit crossing with other species of *Passiflora*.

Although we referred to the partial genomic information of the cultivated passion fruit for enzymatic digestion prediction, the number of SSR markers developed in *Passaflora xishuangbannaensis* is the highest one, indicating that there may be more SSR markers in the Banna passin fruit genome., and the repeat sequencing suggests that its genome is more complex[22].

Ortiz *et al*. [23] found that the transferability of 17 pairs of SSR markers in *Passiflora alata* and *Passiflora edulis* f. *flavicarpa* was 47%. Cazé *et al*. [9]developed 12 pairs of SSR primers in *Passiflora contracta*, but only 2 pairs of markers were successfully amplified in their sister species *Passiflora ovalis*. Silva *et al*. [11]used 31 pairs of SSR primers previously designed by other investigators to study the transferability of SSR markers between wild-type and cultivated passion fruits. The transferability of some SSR markers was very high (71.4%), indicating that it is feasible to establish the transferability SSR marker between wild and cultivated passion fruits.Araya *et al*. [5]analyzed the transferability of the cultivated passion fruit SSR markers using 90 accessions of 78 *Passiflora* species (*Passiflora Ditrophhana, Astrophea* and *Decaloba*), and found that it was 33%-89%. In this study, the average transferability of SSR markers among 10 *Passiflora* accessions was 54.22%, and cultivated passion fruits were higher than that of other species. The transferability of *Passiflora caerulea* and *Passiflora xishuangbannaensis* is extremely low. Therefore, the development of SSR markers in the cultivated passion fruit can increase thetransferability between different *Passiflora* species.

*Passiflora xishuangbannaensis* was currently found only in the Banna region with a wild distribution, and its SSR transferability rate was 0, probably because of the distant relationship between Banna passion fruit and other *Passiflora* accessios. There were large differences in their genome sequences, consistent with the genetic evolution of 10 *Passiflora* accessions. It was also indicated that SSR markers are ideal markers for studying the phylogenetic relationship of *Passiflora* species.

## 5. Conclusions

In this study, we sequenced ten *Passiflora* accessions by RAD-Seq. Using *Passiflora* P4 and P8 as the reference genome, we constructed two phylogenetic trees based on SNPs. The genetic relationship between cultivated passion fruit and other species *Passiflora* is distinct. The SSR transferability in the cultivated passion fruits are higher than that of the other species. The highlights are: (1) successfully finished reduced-representation genome sequencing of *Passiflora* genome by enzyme EcoRI; (2) the first large-scale development of SNP markers in *Passiflora*; (3) developed a large number of cross-species SSR markers; (4) the SSR transferability in cultivated passion fruits is the best.

## Data Availability

The sequencing data were deposited in the Sequence Read Archive (SRA) database under the accession number SRR10272548, SRR10272549, SRR10272550, SRR10272551, SRR10272552, SRR10272553, SRR10272554, SRR10272555, SRR10272556 and SRR10272557.

## Acknowledgments

This work was supported by Guangxi Natural Science Foundation of China (2018GXNSFBA281024), Guangxi’s Ministry of Science and Technology (AB18294007), Guangxi Academy of Agricultural Sciences (2018YT19, TS2016010).

## Author Contributions

Y. Wu., H. Mou and H.Y. contributed to study design, J. Liu. and W. Huang. contributed to data analysis, Q. Tian. contributed to PCR analysis, X. Xia. contributed to primer design, Y. Wu. and H.Y. wrote this manuscript, H. Mou revised the manuscript. All authors read and approve the paper.

